# Baculovirus AC102 is a nucleocapsid protein that is crucial for nuclear actin polymerization and nucleocapsid morphogenesis

**DOI:** 10.1101/248278

**Authors:** Susan E. Hepp, Gina M. Borgo, Simina Ticau, Taro Ohkawa, Matthew D. Welch

**Author notes:** Address correspondence to Matthew D. Welch. Present address: Ohana Biosciences, Cambridge, Massachusetts, USA.

## Abstract

The baculovirus *Autographa californica* multiple nucleopolyhedrovirus (AcMNPV), the type species of alphabaculoviruses, is an enveloped DNA virus that infects lepidopteran insects and is commonly known as a vector for protein expression and cell transduction. AcMNPV belongs to a diverse group of viral and bacterial pathogens that target the host cell actin cytoskeleton during infection. AcMNPV is unusual, however, in that it absolutely requires actin translocation into the nucleus early in infection, and actin polymerization within the nucleus late in infection coincident with viral replication. Of the six viral factors that are sufficient, when coexpressed, to induce the nuclear localization of actin, only AC102 is essential for viral replication and the nuclear accumulation of actin. We therefore sought to better understand the role of AC102 in actin mobilization in the nucleus early and late in infection. Although AC102 was thought to function early in infection, we found that AC102 is predominantly expressed as a late protein. In addition, we observed that AC102 is required for F-actin assembly in the nucleus during late infection, as well as for proper formation of viral replication structures and nucleocapsid morphogenesis. Finally, we found that AC102 is a nucleocapsid protein and a newly recognized member of a complex consisting of the viral proteins EC27, C42, and the actin polymerization protein P78/83. Taken together, our findings suggest that AC102 is necessary for nucleocapsid morphogenesis and actin assembly during late infection through its role as a component of the P78/83-C42-EC27-AC102 protein complex.

**IMPORTANCE:** The baculovirus *Autographa californica* multiple nucleopolyhedrovirus (AcMNPV) is an important biotechnological tool for protein expression and cell transduction, and related nucleopolyhedroviruses are also employed as environmentally benign insecticides. One impact of our work is to better understand the fundamental mechanisms through which AcMNPV exploits the cellular machinery of the host for replication, which may aid in the development of improved baculovirus-based research and industrial tools. Moreover, AcMNPV’s ability to mobilize the host actin cytoskeleton within the cell’s nucleus during infection makes it a powerful cell biological tool. It is becoming increasingly clear that actin plays important roles in the cell’s nucleus, yet the regulation and function of nuclear actin is poorly understood. Our work to better understand how AcMNPV relocalizes and polymerizes actin within the nucleus may reveal fundamental mechanisms that govern nuclear actin regulation and function, even in the absence of viral infection.

## INTRODUCTION

*Autographa californica* multiple nucleopolyhedrovirus (AcMNPV), the type species of alphabaculoviruses, is an enveloped DNA virus that infects the larvae of lepidopteran insects (1). Like other NPVs, it has a circular genome (of 134 kb), replicates in the cell’s nucleus, and produces two different viral forms, a budded virus (BV) that buds from the plasma membrane and an occlusion-derived virus (ODV) that is enveloped in the nucleus and enclosed in large crystalline bodies called polyhedra. AcMNPV is most commonly used as a vector for protein expression and cell transduction (2). It also belongs to a diverse group of viral and bacterial pathogens that target the host cell actin cytoskeleton during infection by inducing the polymerization of actin monomers (G-actin) into actin filaments (F-actin) to enable intracellular actin-based motility (3, 4). AcMNPV is unusual, however, in that it uses host actin in both the cytoplasm and the nucleus (5, 6), and that it absolutely requires actin polymerization for progeny virus production (7–11).

AcMNPV mobilization of host actin begins immediately after enveloped virions fuse with the plasma membrane or endosomal membrane and viral nucleocapsids are released into the host cell cytoplasm. Nucleocapsids then initiate actin-based motility (12) and form actin comet tails (5, 6), which speeds transit to the nucleus (12). Expression of early viral genes induces the translocation of G-actin into the nucleus, a phenomenon referred to as nuclear localization of actin (NLA) (13). Interestingly, NLA can also be induced in uninfected cells by expressing combinations of six viral genes, including the immediate early transcriptional transactivator *ie-1*, as well as *pe38*, *ac004*, *ac152*, and either *ac102* or *he65* (13). Of the latter five genes, only *ac102* is essential for viral replication and the nuclear accumulation of G-actin, indicating it plays a key role in NLA (14).

AcMNPV also mobilizes actin during the late stage of infection (5), when the G-actin that is imported into the nucleus during early infection is polymerized into F-actin in the nuclear ring zone (RZ) (14) surrounding the central virogenic stroma (VS), where DNA replication and nucleocapsid assembly occur. Nuclear F-actin polymerization requires the activity of the viral minor capsid protein P78/83 (encoded by *ac009*), a mimic of host Wiskott-Aldrich Syndrome protein (WASP) family proteins (15) that activates the host Arp2/3 complex to promote actin assembly (9). P78/83 also forms a complex with the viral proteins BV/ODV-C42 (C42; encoded by *ac101*) and ODV-EC27 (EC27; encoded by *ac144*) (16) and associates with one end of the nucleocapsid (17). The genes encoding all three of these proteins are essential for nucleocapsid morphogenesis and the production of budded virus (BV) progeny (9, 18, 19).

To better understand how AcMNPV coordinates actin mobilization early and late in infection, we further investigated the role of NLA factor AC102. We made the surprising observation that AC102 is predominantly expressed as a late gene. In addition, we observed that AC102 is required for F-actin assembly in the nucleus during late infection, as well as for normal VS formation and nucleocapsid morphogenesis. Finally, we found that AC102 is a nucleocapsid protein that it copurifies with EC27, C42, and P78/83. Taken together, our findings suggest that AC102 is necessary for nucleocapsid morphogenesis and P78/83-dependent F-actin assembly during late infection through its role as a component of the P78/83-C42-EC27-AC102 complex.

## RESULTS

### AC102 is predominantly expressed late in infection

Although AC102 was presumed to be an early gene based on its activity in promoting the NLA phenotype, the timing of AC102 expression had not previously been investigated. We therefore began by determining the temporal expression profile of AC102 by Western blotting over a range of times post infection with the wild-type AcMNPV strain WOBpos (9), using a polyclonal antibody we generated that specifically recognizes AC102 (**Fig. 1A**). Surprisingly, at 0, 8, and 10 hpi, no detectable AC102 was present. Expression of AC102 was first detected at 12 hpi and protein continued to accumulate through 36 hpi. The timing of AC102 expression matched that of VP39, the major capsid protein and a tightly regulated late factor (20).

**Fig. 1:**
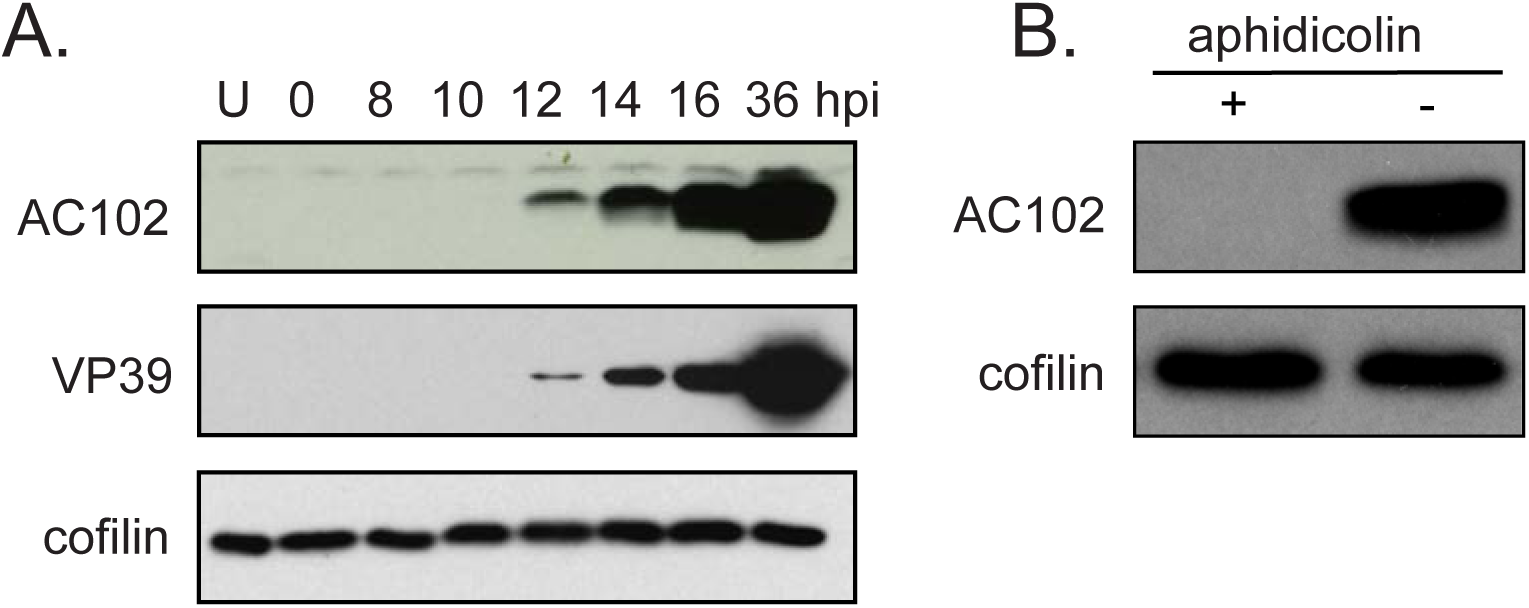
AC102 is predominantly expressed late in infection. **(A)** Western blots of lysates from uninfected Sf9 cells (U), and cells at various time post infection with WOBpos at an MOI of 10, probed for AC102, VP39 (a late protein) and cofilin (loading control). **(B)** Western blots of lysates from WOBpos-infected cells treated with 5 μg/ml aphidicolin (+) or DMSO control (−) at 0 hpi, and then processed at 24 hpi and probed for AC102 and cofilin (a loading control).

To confirm the lack of detectable AC102 expression during the early phase of infection, prior to the initiation of DNA replication, infected cells were treated with aphidicolin, a drug that inhibits DNA synthesis and thus prevents the early to late transition. No AC102 was detected by Western blotting after aphidicolin treatment (**Fig. 1B**). These data indicate that *ac102* is predominantly expressed as a late gene.

### The AC102-K66A mutation results in 10-fold reduced viral titers and small plaques

Given that AC102 is predominantly expressed late in infection, we next sought to investigate the late function(s) of AC102. Because *ac102* is an essential gene (14, 21), it is not possible to produce a virus carrying a deletion mutation. Therefore, we generated ten mutant viruses carrying one of the following point mutations in AC102: N47A, T53A, A55V, D61A, K66A, S77A, A80V, L96A, L105A, or N114A. These amino acid residues were chosen because they are highly conserved between orthologs of AC102 in diverse alphabaculoviruses (**Fig. 2**). Growth of the mutants was initially assessed by infecting cells at MOI of 10 and then measuring viral titer at 18 or 48 hpi (**Table 1**). One virus, containing the AC102-D61A mutation, did not produce any detectable progeny, suggesting this mutation causes inviability. Most other mutants exhibited modest 2- to 5-fold reductions in viral titer compared with WOBpos at one or both time points. The AC102-K66A mutant virus, in contrast, produced 10-fold fewer BV progeny than WOBpos at both time points, and was chosen for further analysis. In a more comprehensive one-step growth curve, the AC102-K66A mutant produced 10-fold fewer progeny at 18, 24, 36, and 48 hpi (**Fig. 3A**, left). To assess whether the AC102-K66A mutation was the cause of the growth defect, an AC102-K66A-rescue virus was generated by inserting a wild-type copy of *ac102* into the bacterial replication cassette in the AC102-K66A bacmid (9). The growth kinetics of the AC102-K66A-rescue virus were indistinguishable from those of WOBpos (**Fig. 3A**, right). Thus, the AC102-K66A mutation causes a 10-fold reduction in BV progeny production throughout the late phase of infection.

**Table 1.** Titers of WOBpos wild type and ten *ac102* mutant viruses at 18 and 48 hpi

| time<sup>a</sup> | WOBpos | *ac102-*<br>*N47A* | *ac102-*<br>*T53A* | *ac102-*<br>*A55V* | *ac102-*<br>*D61A* | *ac102-*<br>*K66A* | *ac102-*<br>*S77A* | *ac102-*<br>*A80V* | *ac102-*<br>*L96A* | *ac102-*<br>*L105A* | *ac102-*<br>*N114A* |
| --- | --- | --- | --- | --- | --- | --- | --- | --- | --- | --- | --- |
| 18 hpi | 3.5x10<sup>5</sup><br>+/- | 1.5x10<sup>5</sup><br>+/- | 3.2x10<sup>5</sup><br>+/- | 1.5x10<sup>5</sup><br>+/- | - | 3.6x10<sup>4</sup><br>+/- | 1.8x10<sup>5</sup><br>+/- | 4.4x10<sup>5</sup><br>+/- | 5.5x10<sup>4</sup><br>+/- | 3.8x10<sup>5</sup><br>+/- | 1.1x10<sup>5</sup><br>+/- |
|  | 0.4x10<sup>5</sup> | 0.4x10<sup>5</sup> | 0.3x10<sup>5</sup> | 0.1x10<sup>5</sup> |  | 0.7x10<sup>4</sup> | 0.1x10<sup>5</sup> | 0.6x10<sup>5</sup> | 1.0x10<sup>4</sup> | 0.5x10<sup>5</sup> | 0.3x10<sup>5</sup> |
| 48 hpi | 7.7x10<sup>7</sup><br>+/- | 4.8 x10<sup>7</sup><br>+/- | 6.3x10<sup>7</sup><br>+/- | 3.9x10<sup>7</sup><br>+/- | - | 7.7x10<sup>6</sup><br>+/- | 7.7x10<sup>7</sup><br>+/- | 1.5x10<sup>7</sup><br>+/- | 6.3x10<sup>7</sup><br>+/- | 3.6x10<sup>7</sup><br>+/- | 2.6 x10<sup>7</sup><br>+/- |
|  | 0.3x10<sup>7</sup> | 0.5x10<sup>7</sup> | 1.3x10<sup>7</sup> | 0.2x10<sup>7</sup> |  | 1.0x10<sup>6</sup> | 0.4x10<sup>7</sup> | 0.2x10<sup>7</sup> | 0.3x10<sup>7</sup> | 0.1x10<sup>7</sup> | 0.2x10<sup>7</sup> |

<sup>a</sup>Cells were infected at an MOI of 10 and progeny virus released into the growth media at 18 and 48 hpi was enumerated by titering. Experiments were done in triplicate and mean titers +/- SD are reported here. *ac102-D61A* produced no viral progeny, and thus could not be amplified for titering.

**Fig. 2:**
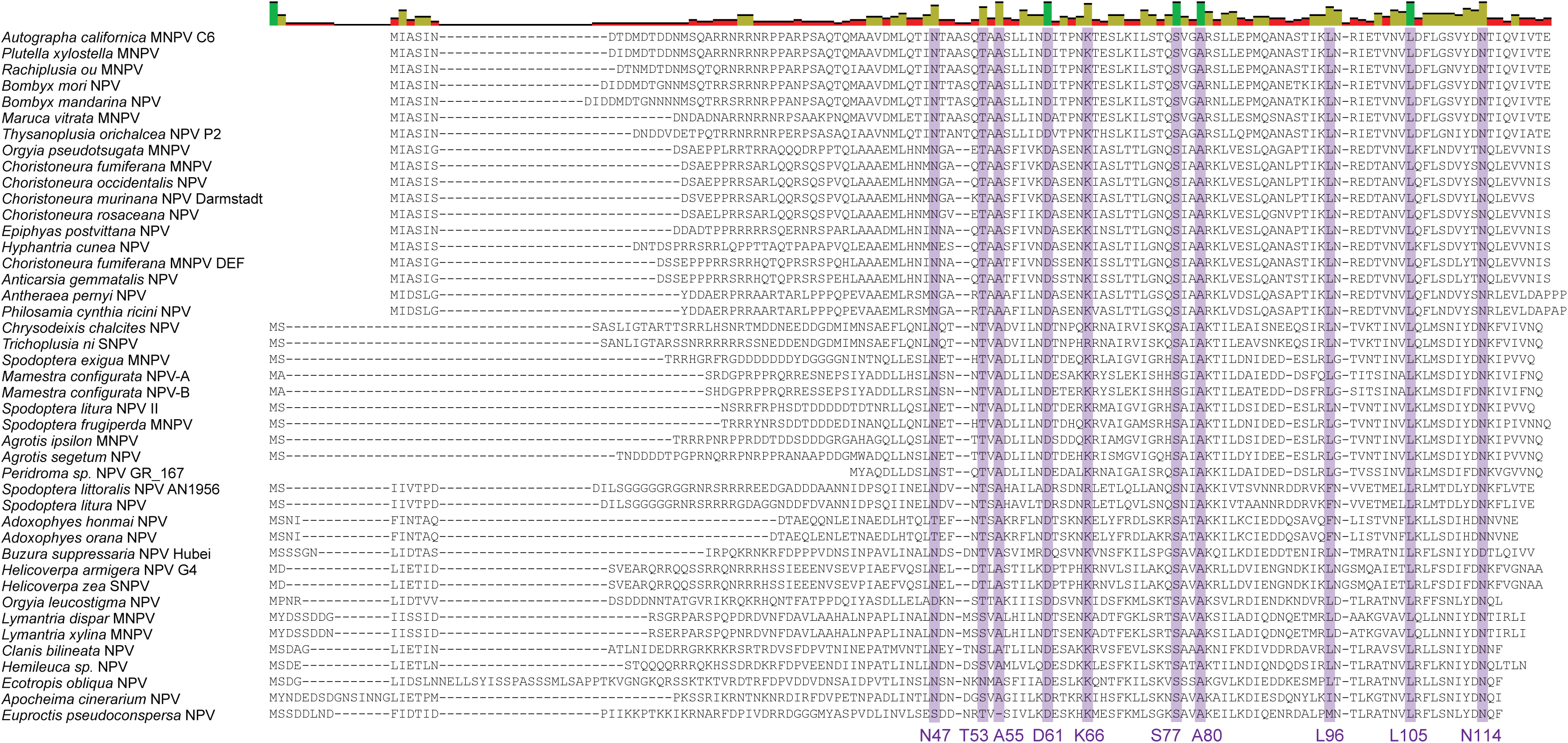
Alignment of AC102 protein sequences from alphabaculoviruses. AC102 amino acid sequences were aligned using MAFFT (56), then manually edited for quality. Residues with at least 85% identity across all alphabaculoviruses are highlighted in purple.

**Fig. 3:**
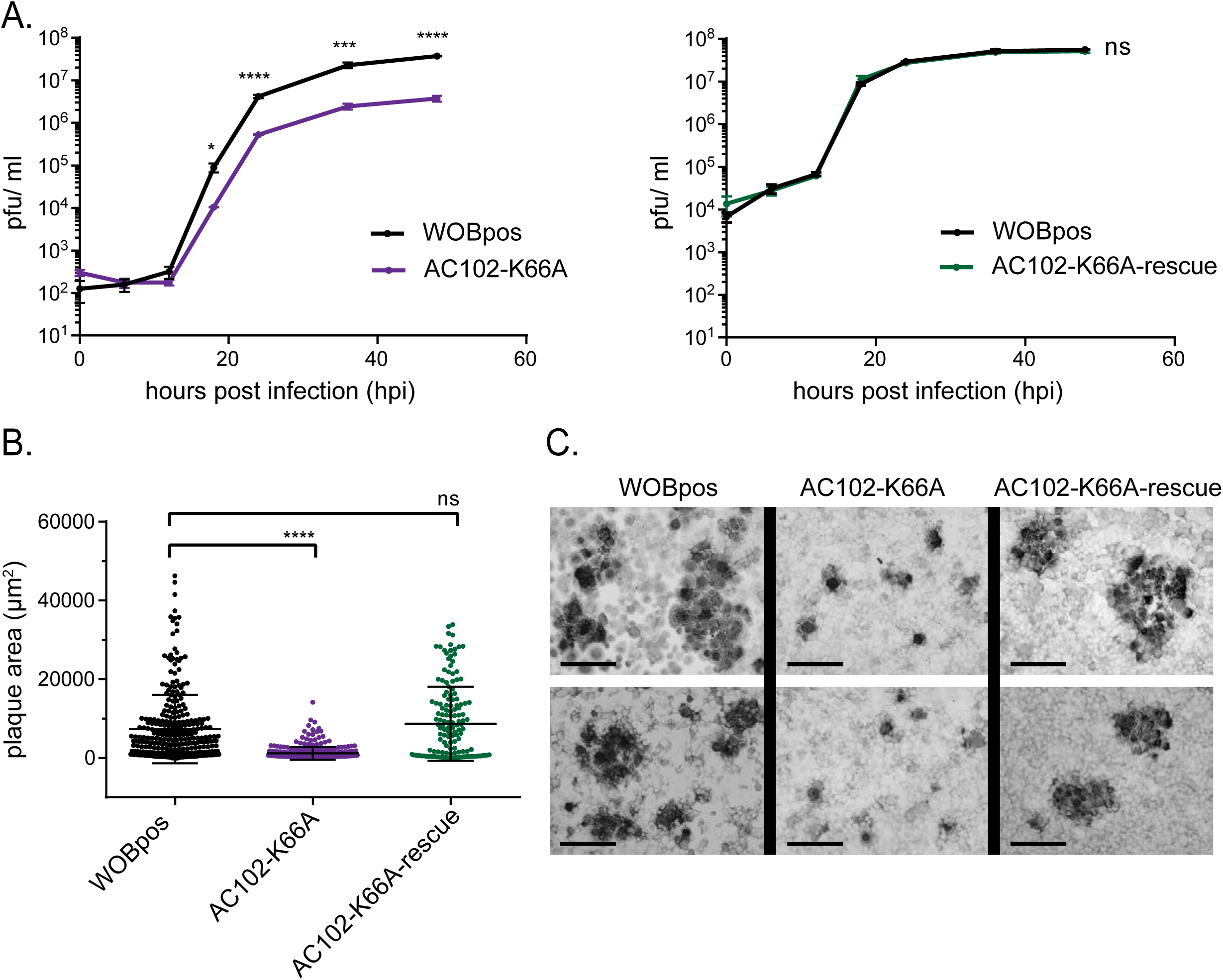
The AC102-K66A mutation results in 10-fold reduced viral titers and a small plaque phenotype. **(A)** One-step growth curves of WOBpos and AC102-K66A viruses (left) or WOBpos and AC102-K66A-rescue viruses (right). Data are the mean +/- SD from three independent experiments. P-values were calculated using a Student’s t-test, and are indicated as follows: ns = non-significant, * = p<0.05, *** = p<0.001, and **** = p<0.0001. **(B)** Quantification of plaque area for WOBpos (n=303), AC102-K66A (n=344), AC102-K66A-rescue (n=160). Each point represents one plaque size measurement, pooled from three independent experiments. Center bar represents the mean, and top and bottom bars represent SD. P-values were calculated by a Kruskal-Wallis test followed by a Dunn’s post test, and are indicated as follows: ns = non-significant, and **** = p<0.0001. **(C)** Plaques resulting from WOBpos, AC102-K66A, or AC102-K66A-rescue viruses visualized by immunostaining for viral envelope protein GP64. Scale bars = 100 μm.

In addition to the replication defect, the AC102-K66A mutant virus produced plaques that were 6-fold smaller than those produced by WOBpos or AC102-K66A-rescue (**Fig. 3B**). These small plaques typically consisted of only one or a few infected cells (**Fig. 3C**). The small plaques suggest that the AC102-K66A mutation results in a reduced capacity to spread from cell to cell.

### The AC102-K66A mutation results in lower AC102 expression

To further investigate the nature of the defect caused by the AC102-K66A mutation, we compared the relative levels of wild-type AC102 and mutant AC102-K66A protein over the course of infection using Western blotting (**Fig. 4A, B**). For both, initial expression was observed at 12 hpi. However, at all time points tested, the levels of AC102-K66A were significantly reduced compared with wild-type AC102 (**Fig. 4B**). These results indicate that the AC102-K66A mutation does not affect the onset of AC102 expression, but results in lower protein levels, likely by impacting the translation or stability of AC102.

**Fig. 4:**
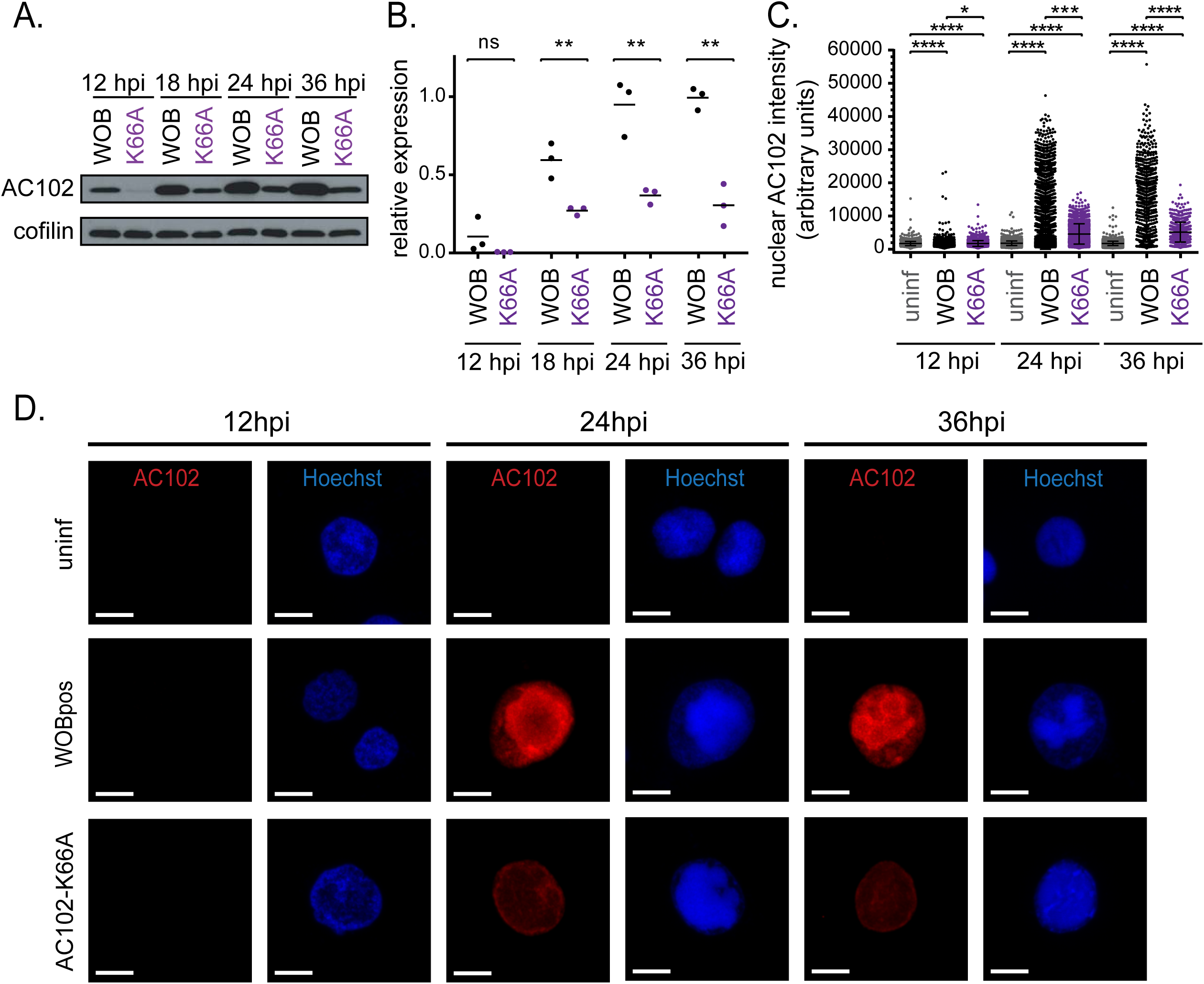
The AC102-K66A mutation results in lower AC102 expression and altered localization to the periphery of the virogenic stroma. **(A)** Western blot of lysates from Sf9 cells infected with WOBpos or AC102-K66A virus at an MOI of 10, prepared at 12, 18, 24, or 36 hpi, and probed for AC102 and cofilin (loading control). **(B)** Quantification of AC102 protein levels from Westerns blots by densitometry. Each dot represents the expression level in one independent experiment relative to the mean for WOBpos at 36 hpi. Lines are the mean from three independent replicates. P-values were calculated using a Student’s t-test and are indicated as follows: ns = non-significant and ** = p<0.01. **(C)** Nuclear intensity of AC102 immunofluorescence staining in uninfected Sf21 cells (uninf), or cells infected with WOBpos or AC102-K66A virus at an MOI of 10, and fixed at 12, 24, or 36 hpi. Bars are the mean from three independent replicates, and error bars are SD. P-values were calculated by the Kruskal-Wallis test followed by a Dunn’s post test and are indicated as follows: ns = non-significant, * = p<0.05, *** = p<0.001, and **** = p<0.0001. **(D)** Sf21 cells stained for AC102 (immunofluorescence; red) and DNA (Hoechst; blue). Scale bars = 10 μm.

We also observed the effect of the AC102-K66A mutation on the abundance and localization of AC102 in infected cells by immunofluorescence microscopy. At 12 hpi, AC102 was not visible in WOBpos or AC102-K66A-infected cells (**Fig. 4C, D**). At 24 and 36 hpi, WOBpos-infected cells showed strong AC102 signal in the nucleus, particularly in and around the VS (identified by a region of intense staining of viral DNA). In contrast, most AC102-K66A-infected cells showed reduced levels of AC102 in the nucleus with staining that was often limited to a thin outline of the VS at 24 and 36 hpi (**Fig. 4C, D**). Thus, the AC102-K66A mutation causes a reduction of AC102 and a redistribution to the periphery of the VS.

### AC102 promotes a condensed virogenic stroma and is important for nucleocapsid assembly

To further elucidate the roles of AC102 in the nucleus, we examined the effect of the AC102-K66A mutation on nuclear organization and nucleocapsid assembly. We initially noted that in AC102-K66A-infected cells, the VS appeared to be less condensed than that of WOBpos-infected cells, with the structure taking up almost the entire nucleus (**Fig. 4D**). To confirm the effect of AC102-K66A on the VS, we performed immunofluorescence microscopy using an antibody against PP31, a delayed early protein and VS marker (22). The timing of PP31 expression appeared similar in WOBpos and AC102-K66A-infected cells (**Fig. 5A, C**). At 12 hpi, only dim PP31 signal could be detected in the nuclei of both AC102-K66A and WOBpos-infected cells (**Fig. 5A**). At 24 and 36 hpi, PP31 was localized to the condensed VS in WOBpos-infected cells, and to a less condensed VS in AC102-K66A-infected cells. At both time points, PP31 was also less abundant in the VS in AC102-K66A-infected cells compared with WOBpos-infected cells. These results confirm that the AC102-K66A mutation results in an aberrant and decondensed VS structure.

**Fig. 5:**
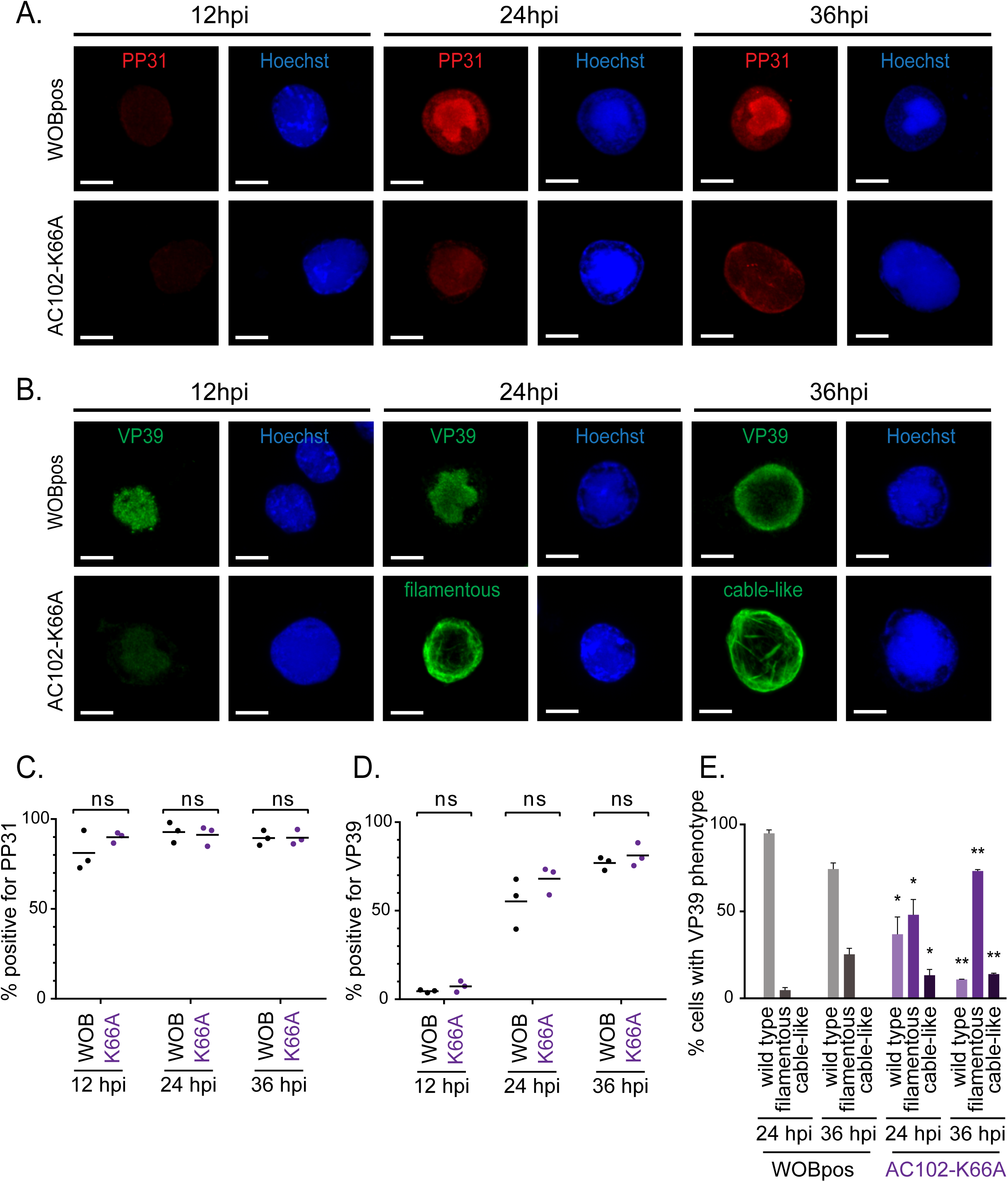
AC102 is important for the maintenance of a condensed virogenic stroma and for normal VP39 localization. **(A)** Sf21 cells infected with WOBpos or AC102-K66A at an MOI of 10, fixed at the indicated times post infection, and stained for PP31 (immunofluorescence; red) and DNA (Hoechst; blue). **(B)** Sf21 cells infected with WOBpos or AC102-K66A at an MOI of 10, fixed at the indicated times, and stained for VP39 (immunofluorescence; green) and DNA (Hoechst; blue). For (A, B) scale bars = 10 μm. **(C, D)** Quantification of the percent cells infected with WOBpos or AC102-K66A, and fixed at the indicated times, that express (C) PP31 or (D) VP39. Each dot represents the average percentage of cells positive for each marker for one independent experiment. Lines are the mean from three independent replicates. Based on a Student’s t-test, there were no significant differences (ns) at each time point tested. **(E)** Quantification of the percentage of cells infected with WOBpos or AC102-K66A, and fixed at the indicated times, that exhibit each VP39 phenotype: “wild type,” with punctate or diffuse VP39 distribution in the VS or RZ; “filamentous,” with long and thin VP39 filaments localized in the RZ; and “cable-like,” with thick VP39 cables localized in the RZ. Data are the mean +/- SD for two independent experiments, with 100-400 cells counted per experiment. P-values were calculated using a Student’s t-test by comparing like phenotypes between WOPpos and AC102-K66A at each time point, and are indicated as follows: * = p<0.05 and ** = p<0.01.

We also investigated the effect of AC102-K66A on the expression and localization of the major capsid protein VP39. As with PP31, the timing of VP39 expression appeared unaffected in the AC102-K66A mutant (**Fig. 5B, D**). At 12 hpi, VP39 showed punctate localization to the VS both in WOBpos and AC102-K66A-infected cells (**Fig. 5B**). At 24 hpi, however, when WOBpos-infected cells continued to show punctate or diffuse VP39 localization in the VS, the majority of AC102-K66A-infected cells had VP39 in the RZ, where it often assembled into long filaments. At 36 hpi, VP39 redistributed to a punctate localization in the RZ in WOBpos-infected cells, whereas it remained filamentous in the ring zone of most AC102-K66A-infected cells. We quantified VP39 distributions in WOBpos and AC102-K66A-infected cells by dividing cells into three phenotypic classes: (1) wild type with punctate distribution in the VS or RZ, (2) “filamentous” with long thin filaments in the RZ, and (3) “cable-like” with thick VP39 cables in the RZ (**Fig. 5E**). At 24 and 36 hpi, the majority of WOBpos-infected cells had wild-type VP39 distribution, whereas the majority of AC102-K66A-infected cells had a filamentous phenotype, and only AC102-K66A-infected cells contained cable-like VP39 structures. Thus, the AC102-K66A mutation causes aberrant assembly of VP39 into filamentous structures late in infection.

We used transmission electron microscopy (TEM) to further compare intranuclear structures in WOBpos and AC102-K66A-infected cells. At 24 hpi, WOBpos-infected cells showed a condensed VS, composed of electron-dense lobes, and a peripheral RZ, both of which contained nucleocapsids (**Fig. 6A, B**). In contrast, AC102-K66A-infected cells lacked a well-defined VS, instead containing a more amorphous region that occupied most of the nucleus and did not contain electron-dense lobes or visible nucleocapsids (**Fig. 6C**). Moreover, in AC102-K66A-infected cells, the peripheral RZ was densely packed with long tubular structures that were variable in length and were often bundled or clustered (**Fig. 6D-F**). The tubules also varied in electron density, with some being nearly electron-lucent and others having electron-dense areas indicating the packaging of viral DNA (**Fig. 6E, F**). Taken together with the VP39 localization studies, these results suggest that the tubular structures are aberrant assemblies of VP39 that are not properly formed into unit-length nucleocapsids, and are instead assembled into long tubular structures that sometimes contain viral DNA.

**Fig. 6:**
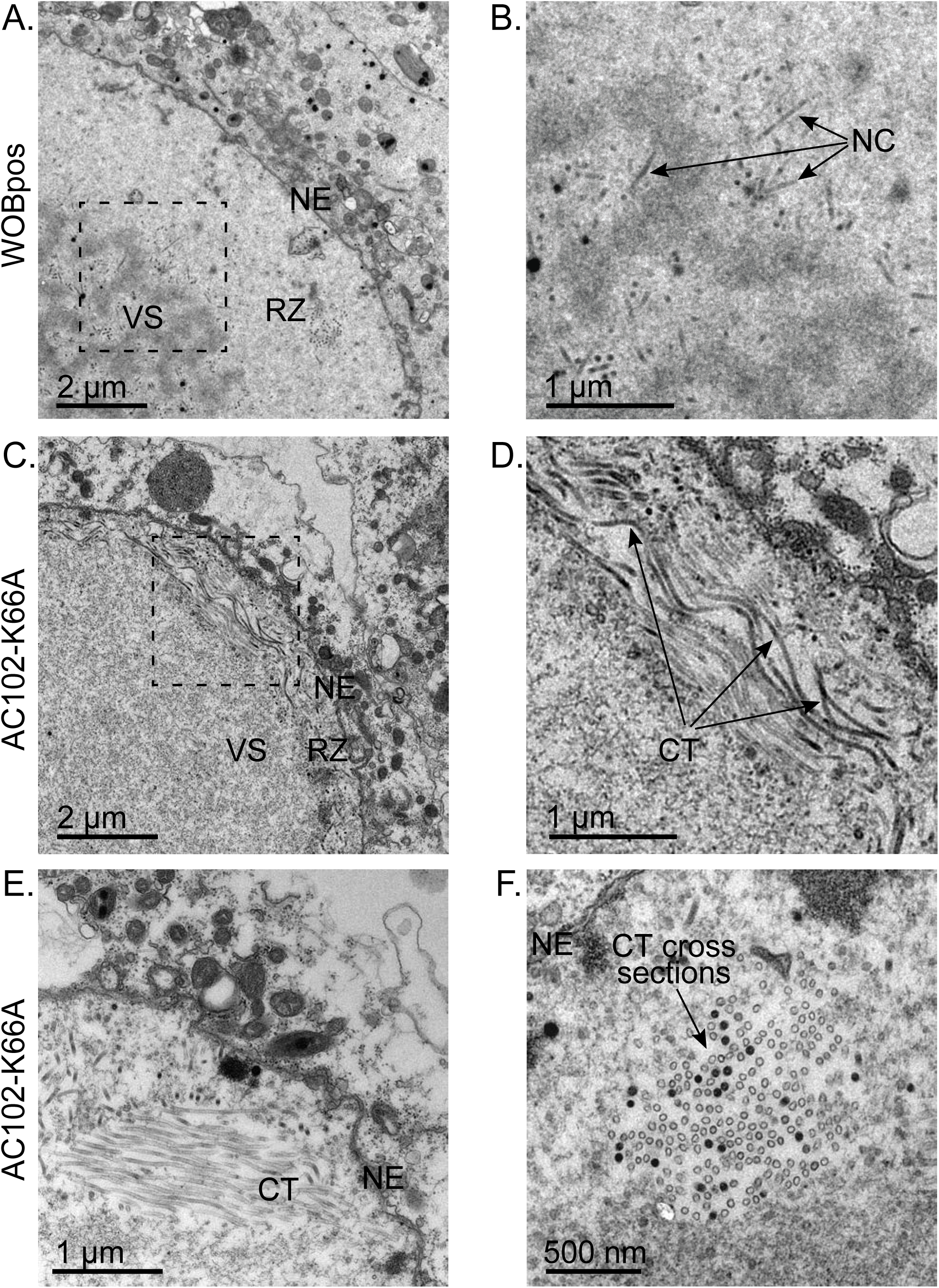
Electron microscopy confirms roles for AC102 in proper nucleocapsid morphogenesis and virogenic stroma formation. **(A)** Electron micrographs of Sf9 cells infected with WOBpos at an MOI of 10, fixed at 24 hpi, and prepared for TEM. The nuclear envelope (NE), ring zone (RZ) and virogenic stroma (VS) are indicated. **(B)** Magnified view of boxed area in (A) showing individual nucleocapsids (NC). **(C)** Electron micrographs of Sf9 cells infected with AC102-K66A, as in (A). The nuclear envelope (NE), ring zone (RZ), and virogenic stroma (VS) are indicated. **(D)** Magnified view of boxed area in (C) showing capsid-like tubule structures (CT). (**E, F**) Electron micrographs of Sf9 cells infected with AC102-K66A, as in (A), showing aggregates of capsid-like tubule structures (CT), in both longitudinal section (E) and cross section (F).

### AC102 is crucial for nuclear actin polymerization in the ring zone late in infection

Our previous work suggested that AC102 is required for nuclear actin polymerization, as transfection of cells with a WOBpos bacmid carrying a deletion of *ac102* (*AcΔ102*) caused a failure in nuclear F-actin accumulation (14). However, the *AcΔ102* virus does not complete a replication cycle, and thus does not necessarily capture the roles of AC102 during the course of a viral infection. To further investigate the role of AC102 in nuclear actin polymerization, we tested the ability of the AC102-K66A virus to cause nuclear accumulation of F-actin late in infection by fluorescence microscopy (**Fig. 7A**). A considerable fraction of WOBpos-infected cells contained nuclear F-actin (defined as a nuclear/cytoplasmic actin intensity ratio of 2 or greater) at 24 and 36 hpi, whereas substantially fewer AC102-K66A-infected cells exhibited nuclear F-actin accumulation at these time points (**Fig. 7B**). These results confirm that AC102 is important for nuclear actin polymerization late in infection.

**Fig. 7:**
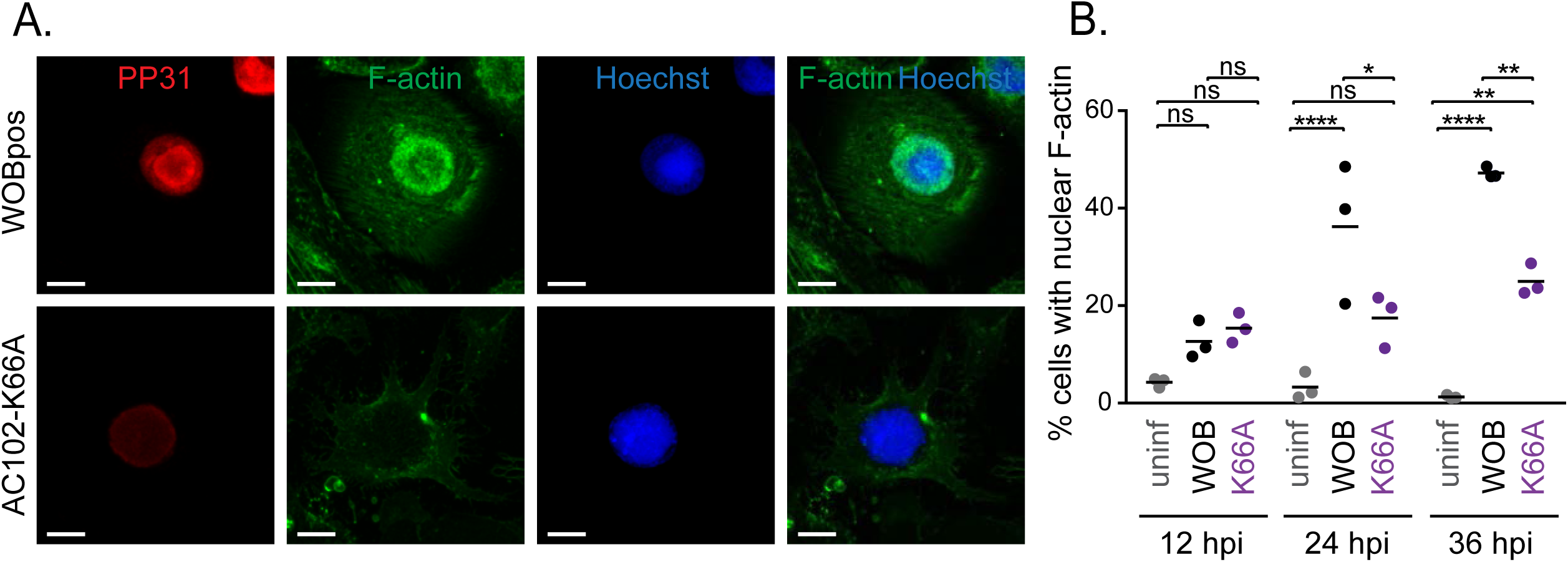
AC102 is crucial for nuclear actin polymerization in the ring zone during late infection. **(A)** Sf21 cells infected with WOBpos or AC102-K66A virus at an MOI of 10, fixed at 36 hpi, and stained for PP31 (immunofluorescence; red), F-actin (Alexa Fluor-phalloidin; green), and DNA (Hoechst; blue). Scale bars = 10 μm. **(B)** Quantification of the percent cells with nuclear F-actin at 12, 24, and 36 hpi, defined as cells with a nuclear to cytoplasmic F-actin intensity ratio of 2 or greater, from Alexa 488-phalloidin staining. Each dot represents the average percentage of cells containing nuclear actin for one independent experiment. Lines indicate the means for three independent experiments. P-values were calculated by one-way ANOVA followed by a Šídák post test and are indicated as follows: ns = non-significant, * = p<0.05, ** = p<0.01, and **** = p<0.0001.

### The AC102-K66A mutation results in a defect in polyhedrin expression and polyhedra formation

We also investigated a possible function for AC102 very late in infection by assessing the timing and expression of the very late protein polyhedrin in WOBpos-infected and AC102-K66A-infected cells by Western blotting (**Fig. 8A, B**). At 18 hpi there was no detectable expression of polyhedrin, as expected for a very late protein. At 24 hpi, WOBpos-infected cells showed strong expression of polyhedrin, whereas AC102-K66A-infected cells showed very little expression. At 36 hpi, polyhedrin expression was significantly higher in WOBpos-infected cells than in AC102-K66A-infected cells. To assess whether lower polyhedrin expression also correlated with fewer polyhedra, we imaged infected cells at 36 hpi using TEM and counted the fraction with at least one polyhedron (**Fig. 8C, D**). Compared with WOBpos-infected cells, significantly fewer AC102-K66A-infected cells contained polyhedra. Together, these data indicate that perturbing AC102 function impacts the proper timing of polyhedrin expression and the formation of polyhedra during the transition to very late stage infection.

**Fig. 8:**
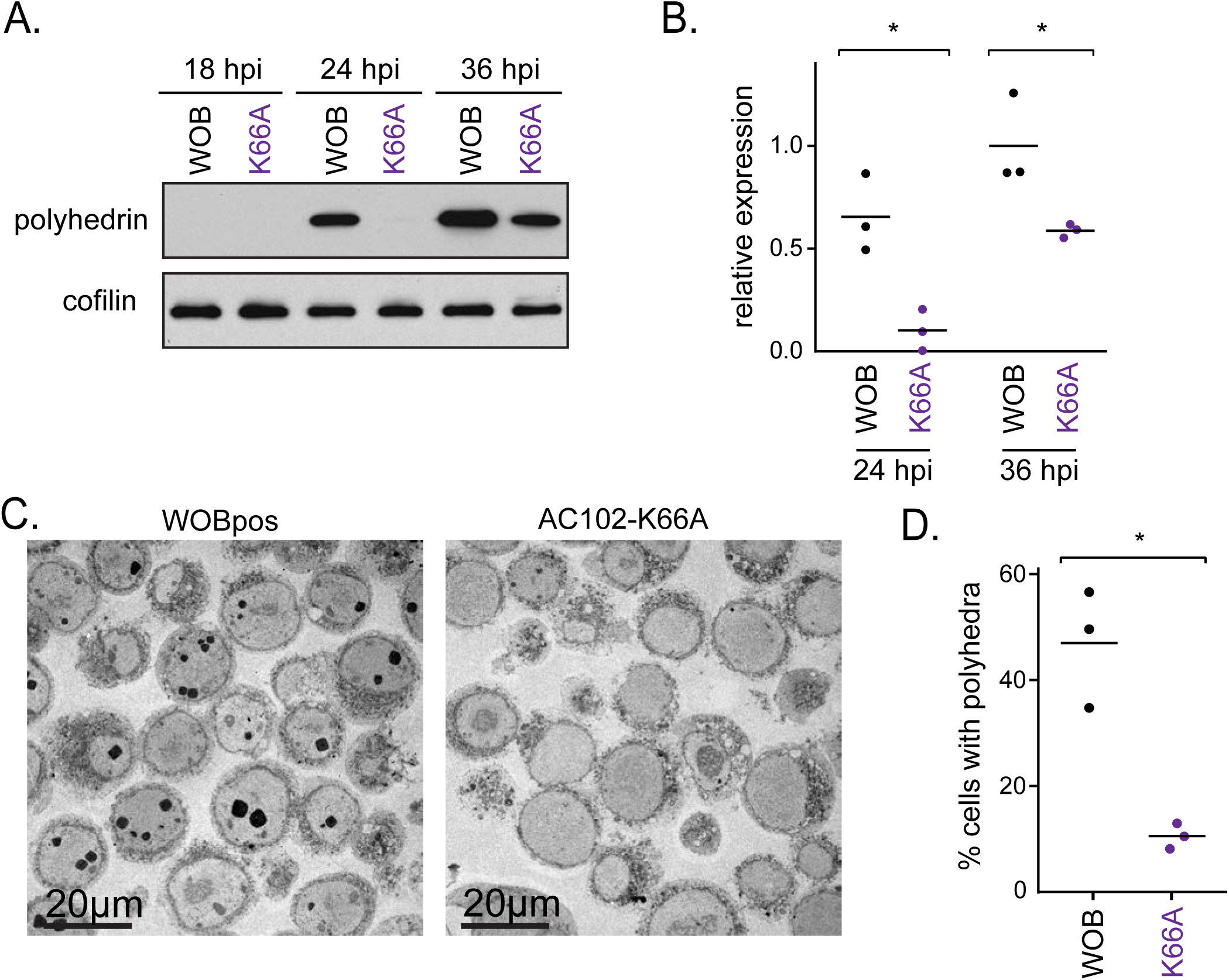
AC102 is important for polyhedrin expression and polyhedra formation very late in infection. **(A)** Western blots of lysates from Sf9 cells infected with WOBpos at an MOI of 10 and probed at 18, 24, or 36 hpi for polyhedrin (top) or cofilin (loading control). **(B)** Quantification of AC102 protein levels from Western blots. Each dot represents the expression level relative to the mean for WOBpos at 36 hpi in one independent experiment. Lines are the mean from three independent replicates. P-values were calculated using a Student’s t-test and are indicated as follows: * = p<0.05 **(C)** TEM micrographs of WOBpos or AC102-K66A-infected cells at 36 hpi showing the presence or absence of electron-dense polyhedra. **(D)** Percentage of cells with at least one polyhedra for WOBpos or AC102-K66A-infected cells visualized as in (C). Each dot represents the average percentage of cells with at least one polyhedra for one independent experiment, and lines indicate the means for three independent experiments. P-values were calculated using a Student’s t-test and are indicates as follows: *=p<0.05.

### AC102 is a nucleocapsid protein that interacts with P78/83, C42, and EC27

The timing of AC102 expression as a late gene and the effect of the AC102-K66A mutation on nucleocapsid morphogenesis suggested that AC102 may be a structural component of the nucleocapsid. To test whether AC102 is a virion-associated protein, BV particles were isolated via ultracentrifugation over a sucrose cushion, then fractionated into their envelope and nucleocapsid components. Western blotting revealed that AC102 is associated with the nucleocapsid fraction, but not with the envelope fraction (**Fig. 9A**). These results indicate that AC102 is a structural component of nucleocapsids.

**Fig. 9:**
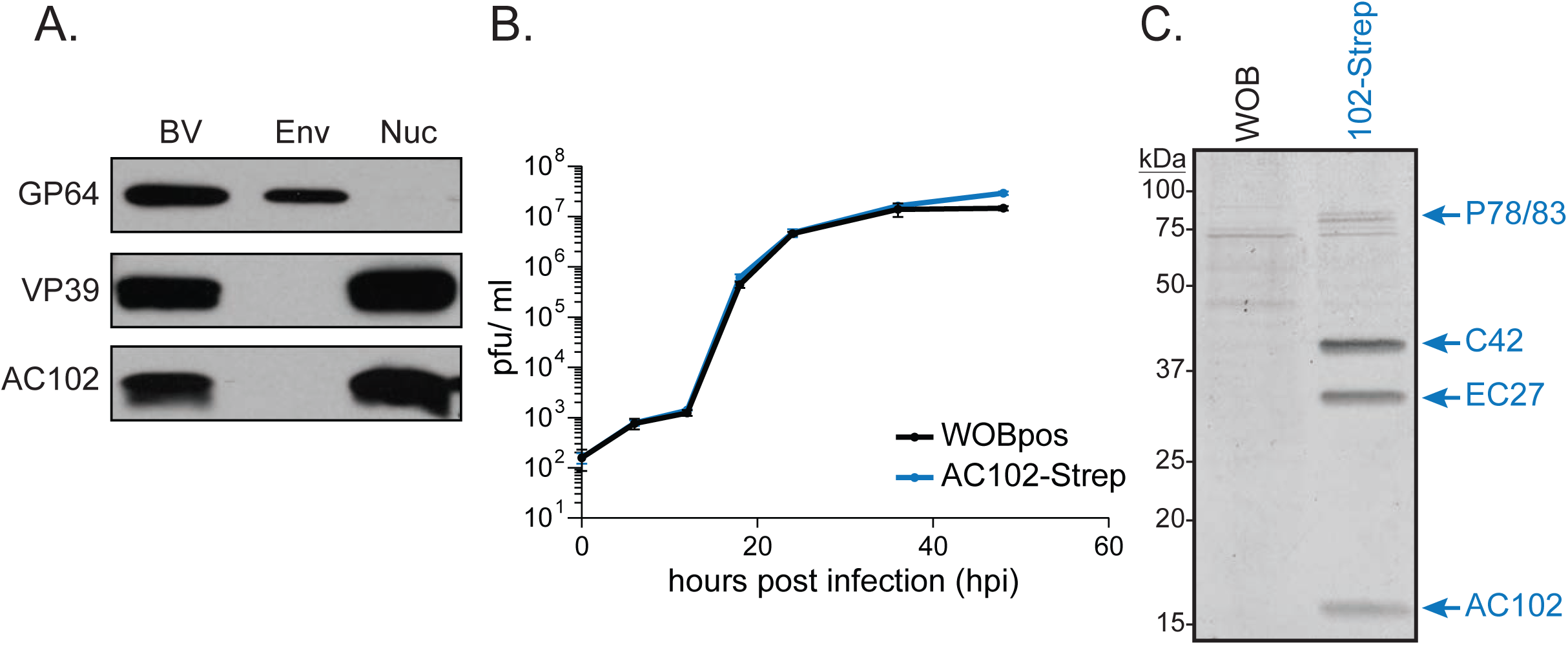
AC102 is a nucleocapsid protein that interacts with P78/83, C42, and EC27. **(A)** Western blots of total budded virus (BV), envelope (Env), and nucleocapsid fractions (Nuc) probed for GP64 (envelope control), VP39 (nucleocapsid control), and AC102. **(B)** One-step growth curves of WOBpos and AC102-Strep viruses in Sf9 cells infected at an MOI of 10. Data are the mean +/- SD from three independent experiments. **(C)** Strep-Tactin-affinity chromatography eluates from WOBpos and AC102-Strep infected Sf9 cell lysates at 24 hpi. Equal amounts of sample were subjected to SDS-PAGE, the gel was stained with Safestain, and bands that were unique to the AC102-Strep sample were excised and analyzed by mass spectrometry. The protein identity of each band is indicated.

As a nucleocapsid structural component, AC102 would be predicted to interact with other capsid proteins. To identify the specific viral proteins with which AC102 interacts, a virus was constructed that expresses AC102 fused to a Twin-Strep-tag (23) in the native *ac102* locus of the viral genome. The AC102-Strep virus grows at a rate that is indistinguishable from WOBpos (**Fig. 9B**), indicating that the AC102-Strep protein is fully functional. To purify AC102 with interacting proteins, Sf9 cells were infected with AC102-Strep or WOBpos as a control, and at 24 hpi cells were lysed and AC102-Strep protein was isolated by Strep-Tactin affinity chromatography. Interestingly, AC102-Strep co-purified with three prominent viral proteins that were identified by SDS-PAGE and mass spectrometry of isolated gel bands as EC27, C42, and P78/83 (**Fig. 9C**), which were previously found to interact with one another in a complex (16). Furthermore, comparative mass spectrometry of AC102-Strep elution samples revealed that AC102 and these three proteins were the most abundant viral proteins in the pulldown (**Table 2**). Other less abundant proteins identified in the pulldown (E25, E18, VP39, and VP80) were also identified as virion-associated proteins by mass spectrometry (24–26). Taken together, these results indicate that AC102 is a nucleocapsid protein that specifically interacts in a complex with the other nucleocapsid proteins P78/83, C42, and EC27.

**Table 2.** Mass spectrometry results from Strept-Tactin-affinity chromatography eluates

| Protein name | Accession | Norm. abundance | Norm. abundance | p-value<sup>b</sup> | HGScore<sup>c</sup> | Protein spectral counts/total spectral counts<sup>d</sup> |  |  |  |
| --- | --- | --- | --- | --- | --- | --- | --- | --- | --- |
|  |  | 102-Strep<sup>a</sup> | control<sup>a</sup> |  |  | 102-Strep 1 | 102-Strep 2 | control 1 | control 2 |
| C42 | P25695 | 3.64 | 0.03 | 0.0002 | 7.417 | 2010/26093 | 3058/30523 | 21/6511 | 26/6785 |
| EC27 | P41702 | 1.94 | 0.03 | 0.0002 | 7.722 | 1019/26093 | 1152/30523 | 15/6511 | 18/6785 |
| P78/83 | Q03209 | 1.10 | 0.03 | 0.0001 | 7.868 | 1227/26093 | 1074/30523 | 28/6511 | 33/6785 |
| AC102 | P41482 | 1.00 | 0.02 | * | N/A | 227/26093 | 244/30523 | 6/6511 | 4/6785 |
| E25 | P41483 | 0.98 | 0.52 | 0.0008 | 5.583 | 345/26093 | 514/30523 | 237/6511 | 225/6785 |
| E18 | P41701 | 0.90 | 0.17 | 0.002 | 4.631 | 78/26093 | 233/30523 | 34/6511 | 24/6785 |
| VP39 | P17499 | 0.55 | 0.09 | 0.006 | 1.888 | 309/26093 | 434/30523 | 65/6511 | 56/6785 |
| VP80 | Q00733 | 0.27 | 0.02 | 0.020 | 0.354 | 334/26093 | 398/30523 | 18/6511 | 26/6785 |

<sup>a</sup>Normalized abundance was calculated by dividing number of spectral counts by protein amino acid length to determine relative abundance, then dividing relative abundance of each protein by the relative abundance of AC102. Positive protein hits are ranked by normalized abundance.
<sup>b</sup>P-values were calculated using the Spotlite web application with the HGScore scoring algorithm.
<sup>c</sup>A higher HGScore means a higher probability of protein-protein interactions.
<sup>d</sup>Numbers are given for two independent replicates from Strep-Tactin affinity eluates from both AC102-Step and control lysates.

## DISCUSSION

AcMNPV protein AC102 was previously shown to be essential for viral replication (14, 21), sufficient to induce nuclear localization of actin (NLA) when expressed with five other viral NLA genes (13), and necessary for NLA (14). Although AC102 was thought to be expressed early in infection (13), here we reveal a role for AC102 as a late-expressed nucleocapsid protein that is a component of the P78/83-C42-EC27-AC102 complex and is crucial for VS organization, nucleocapsid morphogenesis, and nuclear F-actin assembly.

Our findings demonstrate that AC102 is expressed predominantly late in infection, with detectable protein first observed at 12 hpi. The onset of AC102 expression is consistent with the onset of *ac102* mRNA accumulation, which begins at 12 hpi and peaks at 18 hpi based on transcriptomics studies (27). AC102 protein levels continue to accumulate through at least 36 hpi, suggesting that mRNA and protein levels may not be directly correlated at later time points. The onset of AC102 expression is similar to that of VP39, which is strictly expressed late in infection (20). Furthermore, AC102 expression is undetectable after treating cells with aphidicolin, which prevents late gene expression. Thus, *ac102* is predominantly expressed as a late gene.

Many late viral proteins are structural components of virions, and our data indicate that AC102 associates with BV nucleocapsids, but not with their envelopes. Furthermore, we find that AC102 interacts in a stable complex with the nucleocapsid proteins P78/83, C42, and EC27, which were previously shown to interact with one another (16). AC102 had not, however, been recognized as a member of this complex. The associations between P78/83, C42, and EC27 were discovered through yeast-two-hybrid studies, which rely on the interaction of bait and prey proteins fused with domains from a transcription factor. We have found that AC102 is not functional when fused at its N-terminus or C-terminus with larger proteins (unpublished observations), offering a possible explanation as to why interactions with AC102 may have been missed in two-hybrid assays.

A previous analysis of AC102 function relied on the characterization of an *AcΔ102* mutant following bacmid transfection, which does not result in the production of viral progeny (14), thus precluding a thorough assessment of the role of AC102 throughout infection. Our characterization of the AC102-K66A partial loss-of-function mutant virus alleviates this issue and has revealed new roles of AC102 during late infection. We found that AC102 is important for establishing and/or maintaining a condensed VS, as this structure was expanded and amorphous in AC102-K66A-infected cells. This is similar to what has been seen in cells treated with cytochalasin D (10), a drug that inhibits actin polymerization and also induces proteolysis of the genome packaging protein p6.9 (28). AC102 is also important for nucleocapsid morphogenesis, as aberrant capsid-like tubular structures accumulated in the RZ of AC102-K66A-infected cells as visualized by TEM. Similar tubular structures have been observed in cells treated with cytochalasin D (10, 29), as well as in cells transfected/infected with mutant bacmids/viruses carrying deletions of *ac101/c42* and *ac144/ec27* (19), and other genes important for nucleocapsid assembly including *vlf-1/ac77* (30), *49K/ac142* (19), *38K/ac98* (31), *ac54* (32, 33), *ac53* (34), and *pk-1/ac10* (35). These aberrant tubular structures contain the major capsid protein VP39 (19, 29, 31) and are thought to result from perturbed nucleocapsid assembly. The presence of these structures is consistent with our immunofluorescence microscopy images showing VP39 mislocalized into filaments and cables in the RZ of AC102-K66A infected cells. Taken together, these data confirm that AC102 is important for proper organization of viral replication structures and nucleocapsid morphogenesis, most likely due to its role as a structural component of nucleocapsids.

Our observations indicate that, in cells infected with the AC102-K66A mutant, expression of the very late protein polyhedrin is also delayed and reduced, and fewer infected cells contain polyhedra as compared with WOBpos-infected cells. This phenotype is similar to that recently observed for a virus carrying a point mutation in VP39 (VP39-G276S), which exhibits reduced expression of polyhedrin and fewer polyhedra (36). It was suggested that lower polyhedrin expression in the VP39-G276S mutant might result from the sequestration of viral DNA in aberrant tubular nucleocapsid-like structures, and a similar phenomenon may account for the phenotype of the AC102-K66A mutant.

Consistent with observations from cells transfected with an *AcΔ102* mutant bacmid (14), we also observed decreased levels of nuclear F-actin in late stage AC102-K66A-infected cells. AC102’s association with the P78/83-containing complex suggests a mechanism through which AC102 may impact actin polymerization in the nucleus. P78/83 is a viral a mimic of host Wiskott-Aldrich Syndrome protein (WASP) family proteins (15) that activates the host Arp2/3 complex to promote actin polymerization (9). It is required for both actin-based motility during early infection (12) and nuclear F-actin polymerization during late infection (9). The defect in nuclear actin polymerization seen for the AC102-K66A mutant virus could therefore be caused by perturbed stability, localization or actin polymerization activity of the P78/83-C42-EC27-AC102 complex. Consistent with this notion, deletion of *c42/ac101* also causes a reduction in nuclear F-actin (37). Furthermore, because nuclear F-actin is required for progeny production (7–11), the reduced nuclear F-actin accumulation caused by the AC102-K66A mutation may in part account for reduced production of BV.

Interestingly, apart from their late functions, components of the P78/83-C42-EC27-AC102 complex have also been suggested to have functions early in infection. For example, AC102 activity in NLA can occur in the absence of late gene expression and requires prior expression of viral proteins IE1, PE38, and AC152 (13), suggesting that these other NLA factors might regulate AC102 expression or activity early in infection to promote the nuclear localization of G-actin prior to its polymerization into F-actin. Moreover, C42 and EC27 are reported to have other functions, including a role for C42 in the nuclear import of P78/83 (38), and a role for EC27 as a cyclin-like protein that may mediate host cell cycle arrest during infection (39). How early activity might occur remains unclear, as *p78/83*, *c42/ac101*, *ec27/ac144* and *ac102* are predominantly late genes based on transcriptomics data (27). Nevertheless, this same data indicates that low levels of mRNA of all four genes are present 1 h after viral inoculum was added to cells (27), suggesting the possibility of early expression at low levels. Alternatively, the dose of these structural proteins delivered with the initial viral inoculum may be sufficient to carry out early functions.

In closing, our work reveals that AC102 is a central player that links the early nuclear accumulation of G-actin with later nuclear F-actin assembly and nucleocapsid morphogenesis. Future studies into the mechanistic roles of AC102 will reveal how it contributes to the regulation and activity of the P78/83-C42-EC27-AC102 complex, and how it acts in nucleocapsid morphogenesis and the dramatic relocalization and polymerization of actin into the nucleus during baculovirus infection. Given our relatively limited understanding of the normal regulation and function of nuclear actin in uninfected cells, future studies of AC102 will also enhance our understanding of the diverse cellular functions of actin.

## MATERIALS AND METHODS

### Cell lines and viruses

Sf9 cells were maintained in ESF921 media (Expression Systems) at 28°C in shaker flasks. Sf21 cells were maintained in Grace’s insect media (Gemini Bio-Products) with 10% FBS and 0.1% Pluronic F-68 (Invitrogen) at 28°C in shaker flasks. AcMNPV WOBpos (9) was used as the wild-type virus in this study.

### Generation of recombinant viruses

To generate viruses with point mutations in *ac102*, we first generated transfer vectors carrying the AcMNPV viral fragment KpnI-E (2.0 kb KpnI fragment of PstI-C containing *ac102* (13)) and a downstream chloramphenicol resistance (*cat*) cassette for later use in recombinant bacmid selection. These were cloned into the KpnI site of pBSKS+ (Agilent Technologies). The QuikChange II Site-Directed Mutagenesis Kit (Agilent Technologies) was used according to the manufacturer’s protocol to produce ten mutant transfer vectors encoding AC102 with one of the following point mutations: N47A, T53A, A55V, D61A, K66A, S77A, A80V, L96A, L105A, and N114A. To generate mutant viruses, *E. coli* strain GS1783 (40) (provided by Laurent Coscoy, University of California, Berkeley) was transformed with WOBpos viral DNA carrying a deletion of *ac102* (*Ac*Δ*102*) (14). Transformed cells were then shifted to 42°C for 15 min to express recombinase proteins, then subsequently electroporated with linearized mutant transfer vectors. Recombinant bacmids were selected by plating transformed cells on LB containing chloramphenicol and kanamycin. To generate the AC102-K66A-rescue virus, the transfer vector pAC102-rescue was engineered by amplifying *ac102* and its native promoter from viral DNA using PCR, and then subcloning it into pWOBGent3 (12). *E. coli* strain GS1783 containing the AC102-K66A mutant bacmid was shifted to 42°C as described above and then electroporated with linearized pAC102-rescue DNA. Recombinant bacmids were selected by plating transformed cells on LB containing gentamycin. This resulted in one copy of *ac102* under the control of its native promoter being inserted into the AC102-K66A bacmid just upstream of the kanamycin resistance cassette. To generate the AC102-Strep virus, the transfer vector pAC102-StrepII was engineered by amplifying a gene encoding AC102 tagged with a Twin-Strep-tag (23). *E. coli* strain GS1783 containing the *Ac*Δ*102* bacmid (14) was shifted to 42°C as described above, then electroporated with linearized pAC102-StrepII DNA. This resulted in the insertion of the *ac102-Strep* gene into the native *ac102* locus. Recombinant bacmids were selected for by plating transformed *E. coli* onto LB plates containing chloramphenicol. In all cases, isolated bacmid DNA was transfected into Sf9 cells using TransIT-Insect Transfection Reagent (Mirus Bio LLC), and virus was recovered from transfected cell supernatants. For all recombinant viruses, we confirmed proper homologous recombination and the presence of *ac102* point mutations by restriction endonuclease digestion analysis of viral DNA, as well as by PCR and DNA sequencing.

### Viral growth curves and plaque size measurements

To compare the kinetics of progeny BV production for WOBpos, AC102-K66A, AC102-K66A-rescue, and AC102-Strep viruses, one-step growth curves were performed in triplicate using an immunoplaque assay (41). To measure plaque size, images of plaques were captured on an Olympus IX71 microscope with a 20x objective (Olympus LUCPlanFL, 0.45 NA), CoolSNAP HQ camera (Photometrics), and μManager software (42). The Fiji distribution of ImageJ (43) was used to measure the area of individual plaques.

### Budded virus purification and fractionation

To obtain purified BV, Sf9 cells were infected with WOBpos at an MOI of 10 and cell culture supernatant containing BV was collected at 2 dpi. The supernatant was then overlaid onto a 40% sucrose cushion and centrifuged at 100,000 × *g* for 1 h at 15°C using a Beckman SW-28 rotor in a Beckman L8-M ultracentrifuge to pellet BV particles. BV was then resuspended in 10 mM Tris buffer (pH 8.5). To further fractionate BV into envelope and nucleocapsid components, NP-40 was added to 100 μg of BV at a final concentration of 1%, incubated for 1 h at 4°C, and the sample was centrifuged at 80,000 × *g* for 1 h at 15°C using a Beckman TLA-100 rotor in a Beckman TL-100 ultracentrifuge to separate the supernatant fraction containing viral envelope proteins from the pellet fraction containing nucleocapsid-associated proteins. The nucleocapsid fraction was then washed and centrifuged a second time under identical conditions to ensure that there was no contamination from the envelope fraction.

### Purification and identification of AC102 interacting proteins

To purify AC102-Strep together with interacting proteins, Sf9 cells were infected with AC102-Strep or control WOBpos viruses at an MOI of 10, and infected cells were harvested at 24 hpi. Cells were lysed on ice for 10 min in lysis buffer (50 mM Tris-HCl, 150 mM NaCl, 1 mM EDTA, 1% Triton X-100, 1 μg/ml LPC, 1 μg/ml aprotinin, and 1 mM PMSF) and centrifuged in a microcentrifuge at 16,000 × *g* for 2 min at room temperature to separate nuclei and cellular debris from the cytoplasmic supernatant. The clarified cell lysate was incubated with 50 μl (packed volume) of Strep-Tactin Sepharose resin (IBA Lifesciences) for 2 h at 4°C with rotation. Beads were washed with lysis buffer containing 300 mM NaCl, and protein was eluted with lysis buffer containing 6 μM D-desthiobiotin (IBA Lifesciences). Protein concentration was assessed via Bradford assay. Samples containing equal amounts of protein were separated by SDS-PAGE, and gels were stained with SimplyBlue SafeStain (Thermo Fisher Scientific) according to the manufacturer’s maximum sensitivity protocol.

For identification of individual proteins visualized by SDS-PAGE, bands that were unique to the AC102-Strep pulldown were cut out, in-gel digested with trypsin (44), and subjected to mass spectrometry as described below. In addition, for the bulk identification of eluted proteins, whole elution samples from three independent affinity chromatography experiments for both AC102-Strep and control WOBpos were trypsin digested and subjected to mass spectrometry as described below.

Mass spectrometry was performed by the Vincent J. Coates Proteomics/Mass Spectrometry Laboratory at the University of California, Berkeley. For whole elution samples, a nano LC column consisting of 10 cm of Polaris c18 5 μm (Agilent Technologies), followed by 4 cm of Partisphere 5 SCX (GE Healthcare Life Sciences), was washed extensively with buffer A (5% acetonitrile/ 0.02% heptaflurobutyric acid (HBFA)). The column was then directly coupled to an electrospray ionization source mounted on a Thermo-Fisher LTQ XL linear ion trap mass spectrometer. An Agilent 1200 HPLC delivering a flow rate of 300 nl/min was used for chromatography. Peptides were eluted using an 8-step MudPIT procedure (45). The buffers used were: buffer A (above); buffer B (80% acetonitrile/ 0.02% HBFA); buffer C (250 mM ammonium acetate/ 5% acetonitrile/ 0.02% HBFA); and buffer D (500 mM ammonium acetate/ 5% acetonitrile/ 0.02% HBFA). This protocol was also followed for proteins originating from gel bands, except that the nano LC column was packed with 10 cm of Polaris c18 5 μm only, and the chromatography consisted of a simple gradient from 100% buffer A to 40% buffer A, 60% buffer B.

Protein identification and quantification were done with Integrated Proteomics Pipeline IP2 software (Integrated Proteomics Applications) using ProLuCID/Sequest (46), DTASelect (47, 48), and Census (49). Tandem mass spectra were extracted into ms1 and ms2 files from raw files using RawExtractor (50). Data was searched against the AcMNPV (NC_001623.1) translated protein database, supplemented with sequences of common contaminants, and concatenated to a decoy database in which the sequence for each entry in the original database was reversed (51). LTQ data was searched with 3000.0 milli-amu precursor tolerance, and the fragment ions were restricted to a 600.0 ppm tolerance. All searches were parallelized and searched on the VJC proteomics cluster. Search space included all fully tryptic peptide candidates with no missed cleavage restrictions. Carbamidomethylation (+57.02146) of cysteine was considered a static modification. We required 1 peptide per protein and both trypitic termini for each peptide identification. The ProLuCID search results were assembled and filtered using the DTASelect program with a peptide false discovery rate (FDR) of 0.001 for single peptides and a FDR of 0.005 for additional peptides for the same protein. The estimated false discovery rate was about 1% for the datasets used. To better distinguish direct from indirect interactions in whole elution samples, we used Spotlite (52), a web-based platform designed to identify specific protein-protein interactions from affinity purified samples subjected to mass spectrometry. We used the HGSCore scoring algorithm (53) within Spotlite to identify high confidence interactions with a p-value of p < 0.05.

### AC102 purification and anti-AC102 antibody generation

To express recombinant AC102 protein, *ac102* was amplified by PCR and subcloned into the SspI site of pET-1M (provided by the University of California, Berkeley, QB3 MacroLab) to generate plasmid pET-1M-AC102 that encodes a fusion protein of AC102 with an N-terminal 6xHis tag, maltose-binding protein, and tobacco-etch virus protease cleavage site (His-MBP-TEV-AC102). *E. coli* strain BL21(DE3) (New England Biolabs) was transformed with pET-1M-AC102, grown at 37°C to an OD of 0.6-0.8, induced with 250 μM ITPG for 2 h, and then harvested. *E. coli* were resuspended in lysis buffer (100 mM Tris-HCl, pH 8.0, 150 mM NaCl, 1 mM EDTA, 1μg/ml LPC, 1μg/ml aprotinin, and 1mM PMSF), lysed by sonication, and lysates were centrifuged at 20,000 × *g* for 20 min at 4°C using an SS34 rotor in a Sorval RC5C Plus centrifuge. Clarified lysates were incubated with 10 ml (packed volume) of amylose resin (New England Biolabs) for 2 h at 4°C with rotation, washed with 10 volumes of column buffer (100 mM Tris-HCl, pH 8.0, 300 mM NaCl, 1 mM EDTA), then eluted with column buffer containing 10 mM maltose. Protein containing fractions were pooled and MBP was cleaved from AC102 using 1 mg/ml TEV Protease. Released His-MBP and uncleaved His-MBP-TEV-AC102 were removed by binding to HisPur Ni-NTA Resin (Thermo Fisher Scientific). Purified AC102 was concentrated to 1 mg/ml with a 3 kDa molecular weight cut-off protein concentrator (Thermo Fisher Scientific).

To generate custom antibodies that recognize AC102, rabbits were immunized with the purified AC102 protein by Pocono Rabbit Farm and Laboratory, using their standard 91-day protocol. For antibody affinity purification, purified and concentrated AC102 was further purified via ion exchange chromatography, using a HiTrap SP HP 1 ml column (GE Healthcare Life Sciences) and coupled to NHS-activated Sepharose 4 Fast Flow resin (GE Healthcare Life Sciences). Anti-AC102 serum was passed over the AC102 affinity resin, and antibodies were eluted with 100 mM glycine pH 2.5. Antibodies were immediately neutralized to pH 7.5 by adding 1 M Tris pH 8.8 and stored at −20°C or −80°C in 50% glycerol.

### Analysis of AC102 and AC102-K66A expression by Western blotting

To observe AC102 protein expression over the course of infection, Sf9 cells were infected in triplicate with WOBpos at an MOI of 10 and infected cells were harvested at 0, 8, 10, 12, 14, 16, and 36 hpi. Cells were lysed on ice for 10 min in lysis buffer (50 mM Tris-HCl, pH 8.0, 150 mM NaCl, 1 mM EDTA, 1% Triton X-100, 1 μg/ml LPC, 1 μg/ml aprotinin, and 1 mM PMSF) and centrifuged in a microcentrifuge at 16,000 × *g* for 2 min at room temperature to separate nuclei and cellular debris from the cytoplasmic supernatant. Cell lysates were subjected to SDS-PAGE, with equal loading of total protein in all lanes (as determined by Bradford assay). Proteins were transferred to nitrocellulose membranes, and probed by Western blotting using: rabbit anti-AC102; mouse-anti-VP39 antibody P10C6 (54) (kindly provided by JaRue Manning); and rabbit-anti-cofilin antibody GA15 as a loading control (kindly provided by Michael Goldberg and Kris Gunsalus). To assess whether AC102 is expressed late in infection following the onset of DNA replication, cells were infected as above in the presence of either 5 μg/ml aphidicolin or DMSO (as a control). At 24 hpi, cells were lysed and subjected to SDS-PAGE and Western blotting as described above using rabbit-anti-AC102 and rabbit-anti-cofilin antibodies.

To compare expression levels of AC102 and other proteins in WOBpos and AC102-K66A-infected cells, Sf9 cells were infected in triplicate as described above, and lysed at 0, 6, 12, 18, 24, and 36 hpi. Cell lysates were prepared and subjected to SDS-Page and Western blotting as described above using: anti-AC102; rabbit-anti-polyhedrin (55) (generously provided by Loy Volkman); and anti-cofilin.

### Immunofluorescence microscopy

For immunofluorescence microscopy, Sf21 cells were seeded onto CELLSTAR black-walled 96-well plates with micro-clear bottoms (Grenier Bio-One) to ~75% confluency and infected in triplicate with WOBpos or AC102-K66A virus at an MOI of 10. At 12, 24, or 36 hpi, cells were fixed with 4% paraformaldehyde in PHEM buffer (60 mM PIPES, pH6.9, 25 mM HEPES, 10 mM EGTA, 2 mM MgCl_2_), permeabilized in PHEM with 0.15% Triton X-100, blocked in PHEM with 5% normal goat serum (MP Biomedicals) + 1% BSA, and processed for immunofluorescence staining as described previously (14). Primary antibodies used were: anti-AC102, 1:500 dilution in PHEM buffer; anti-VP39 P10C6, 1:200; or rabbit-anti-PP31, 1:200 (22) (kindly provided by Linda Guarino). Phalloidin conjugated to Alexa Fluor 488 or 568 at a 1:400 dilution in PHEM buffer (Molecular Probes) was used to visualize F-actin and 500 μg/ml Hoechst was used to visualize DNA.

Cells were imaged with an Opera Phenix High Content Screening System (PerkinElmer) using its confocal spinning disk mode with either 20x water immersion objective (PerkinElmer, NA 1.0, WD 1.7 mm) or 63x water immersion objective (PerkinElmer, NA 1.15, WD 0.6 mm) and the system’s two sCMOS cameras (4.4-megapixel 2100×2100, 16-bit resolution, 6.5μm pixel size). Image analysis was carried out using Harmony High-Content Imaging and Analysis Software (PerkinElmer) on 16-bit images. All analysis was done using maximum intensity projections except for nuclear and cytoplasmic actin quantification, which was done on single z-plane images through the center of cell nuclei.

### Electron microscopy

Sf9 cells were infected in triplicate with WOBpos or AC102-K66A virus at an MOI of 10 and infected cells were fixed at 18 and 36 hpi for 45 min with 1.5% paraformaldehyde and 2.0% glutaraldehyde in 0.05 M sodium cacodylate buffer, pH 7.3. Cell pellets were embedded in 2% agarose, rinsed three times in 0.05 M sodium cacodylate buffer, pH 7.3, and postfixed in a solution of 1% osmium tetroxide, 1.6% potassium ferricyanide, and 0.1 M sodium cacodylate buffer, pH 7.2. Postfixed samples were then embedded in resin, sectioned, and stained with 2% uranyl acetate in 70% methanol, rinsed in decreasing concentrations of methanol, and stained with Reynolds lead citrate. Samples were imaged with a FEI Tecnai 12 transmission electron microscope (Thermo Fisher Scientific) equipped with a 3072 × 3072 pixel Rio 9 CMOS camera (Gatan).

## ACKNOWLEDGEMENTS

We thank Loy Volkman for the rabbit-anti-polyhedrin antibody, Linda Guarino for the rabbit-anti-PP31 antibody, Michael Goldberg and Kris Gunsalus for the rabbit-anti-cofilin GA15 antibody, and JaRue Manning for the mouse-anti-VP39 P10C6 antibody. We thank the following for facilities and technical support: Kent McDonald and Reena Zalpuri from the Robert D. Ogg Electron Microscope Lab, Lori Kohlstaedt from the Vincent J. Coates Proteomics/Mass Spectrometry Laboratory, and Mary West from the QB3 High-Throughput Screening Facility. We are grateful to Loy Volkman for her advice, support, helpful suggestions, and comments on the manuscript. This work was supported by a Hellman Graduate Award to S.H. and by NIH/NIGMS grant GM059609 to M.W.

